# Integrative epigenomics, transcriptomics and proteomics of patient chondrocytes reveal genes and pathways involved in osteoarthritis

**DOI:** 10.1101/038067

**Authors:** Julia Steinberg, Graham R. S. Ritchie, Theodoros I. Roumeliotis, Raveen L. Jayasuriya, Roger A. Brooks, Abbie L. A. Binch, Karan M. Shah, Rachael Coyle, Mercedes Pardo, Christine L. Le Maitre, Yolande F. M. Ramos, Rob G. H. H. Nelissen, Ingrid Meulenbelt, Andrew W. McCaskie, Jyoti S. Choudhary, J. Mark Wilkinson, Eleftheria Zeggini

**Author notes:** Joint first authors (Julia Steinberg, Graham R. S. Ritchie).

## Abstract

**Background:** Osteoarthritis (OA) is a common disease characterized by cartilage degeneration and joint remodeling. The underlying molecular changes underpinning disease progression are incompletely understood, but can be characterized using recent advances in genomics technologies, as the relevant tissue is readily accessible at joint replacement surgery. Here we investigate genes and pathways that mark OA progression, combining genome-wide DNA methylation, RNA sequencing and quantitative proteomics in isolated primary chondrocytes from matched intact and degraded articular cartilage samples across twelve patients with OA undergoing knee replacement surgery.

**Results:** We identify 49 genes differentially regulated between intact and degraded cartilage at multiple omics levels, 16 of which have not previously been implicated in OA progression. Using independent replication datasets, we replicate statistically significant signals and show that the direction of change is consistent for over 90% of differentially expressed genes and differentially methylated CpG probes. Three genes are differentially regulated across all 3 omics levels: *AQP1*, *COL1A1* and *CLEC3B*, and all three have evidence implicating them in OA through animal or cellular model studies. Integrated pathway analysis implicates the involvement of extracellular matrix degradation, collagen catabolism and angiogenesis in disease progression. All data from these experiments are freely available as a resource for the scientific community.

**Conclusions:** This work provides a first integrated view of the molecular landscape of human primary chondrocytes and identifies key molecular players in OA progression that replicate across independent datasets, with evidence for translational potential.

## BACKGROUND

Osteoarthritis (OA) affects over 40% of individuals over the age of 70 [1], and is a leading cause of pain and loss of physical function [2]. There is no curative therapy for OA; instead, disease management targets symptom control until disease progression culminates in joint replacement surgery. OA is a complex disease, with both heritable and environmental factors contributing to susceptibility [3]. Despite the increasing prevalence, morbidity, and economic impact of the disease [1, 2, 4], the underlying molecular mechanisms of OA progression remain poorly characterized.

Emergent high-throughput technologies and bioinformatics analyses of clinical tissues offer the promise of novel functional approaches to disease characterization, as well as biomarker and therapeutic target discovery. Cartilage degeneration is a key feature of OA, thought to be brought about by an imbalance between anabolic and catabolic processes through a complex network of proteins including proteases and cytokines, reviewed in [5-7]. While disease tissues are inaccessible for many other common complex diseases, cartilage is a relevant disease tissue for OA and is readily accessible at joint replacement surgery. This provides an opportunity to deploy multi-omics (DNA CpG methylation, gene expression, and proteomics assays) in order to characterize the molecular processes underpinning disease development in the right tissue, both to fill a gap in our fundamental understanding of disease biology and to identify novel therapeutic opportunities. In recent years, studies examining individual–omics levels have expanded our understanding of OA pathogenesis, reviewed in [8-11]. Here we report the first application of integrated multi-omics across DNA methylation, RNA sequencing and quantitative proteomics from knee joint tissue to obtain a comprehensive molecular portrait of cartilage degeneration in OA patients (Figure 1a). The fundamental question here addresses the biological processes underpinning disease progression within the OA joint, which is of direct clinical relevance to patients suffering from OA. To achieve this, we collected individually-matched pairs of cartilage tissue from patients undergoing joint replacement surgery, with one sample demonstrating advanced degenerative change and the other demonstrating little or no evidence of cartilage degeneration. The findings were then replicated in independent populations of patients undergoing joint replacement. Notably, no functional genomics study carried out to date in knee OA chondrocytes has focused on all three ‐omics levels examined here. Our data highlight disease processes with involvement across multiple levels, and reveal novel and robustly replicating molecular players with translational potential.

**Figure 1.**
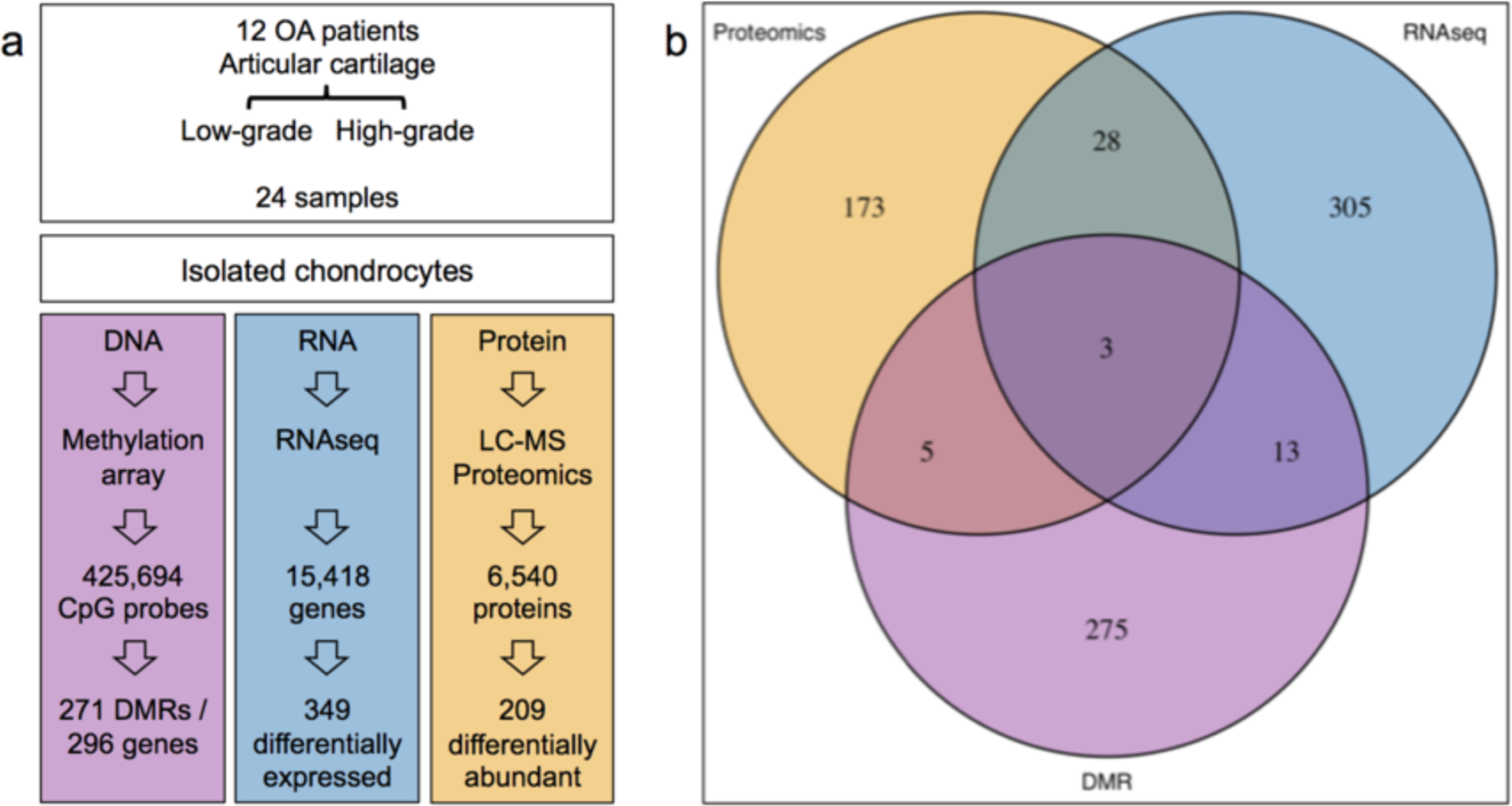
Overview of the genes identified in each ‐omics experiment and their overlap. a) Schematic view of the 3 functional genomics experiments identifying the number of genes shortlisted for each. b) Venn diagram identifying the number of overlapping shortlisted genes from each individual experiment.

## RESULTS

In the discovery population we extracted cartilage and subsequently isolated chondrocytes from the knee joints of 12 OA patients undergoing total knee replacement surgery (see **Methods**). We obtained two cartilage samples from each patient, scored using the OARSI cartilage classification system [12, 13]: one sample with high OARSI grade signifying highgrade degeneration (referred to as “degraded sample”), and one sample with low OARSI grade signifying healthy tissue or low-grade degeneration (referred to as “intact sample”) (Supplementary Figure 1). We compared the degraded and intact tissue across patient-matched samples.

### Quantitative proteomics

We used isobaric labeling liquid chromatography-mass spectrometry (LC-MS) to quantify the relative abundance of 6540 proteins that mapped to unique genes. This is the most comprehensive differential proteomics study on OA samples to date. We identified 209 proteins with evidence of differential abundance (Supplementary Table 1); ninety were found at higher abundance in the degraded samples, and 119 were found at lower abundance. For two representative patients we also used an orthogonal label-free approach that confirmed the protein quantification data (Supplementary Figure 2). For three of the differentially abundant proteins (ANPEP, AQP1 and TGFBI), we validated the higher levels in degraded cartilage by Western blotting (Supplementary Figure 3). All three were also detected as significant in the RNA sequencing data (see below).

**Figure 2:**
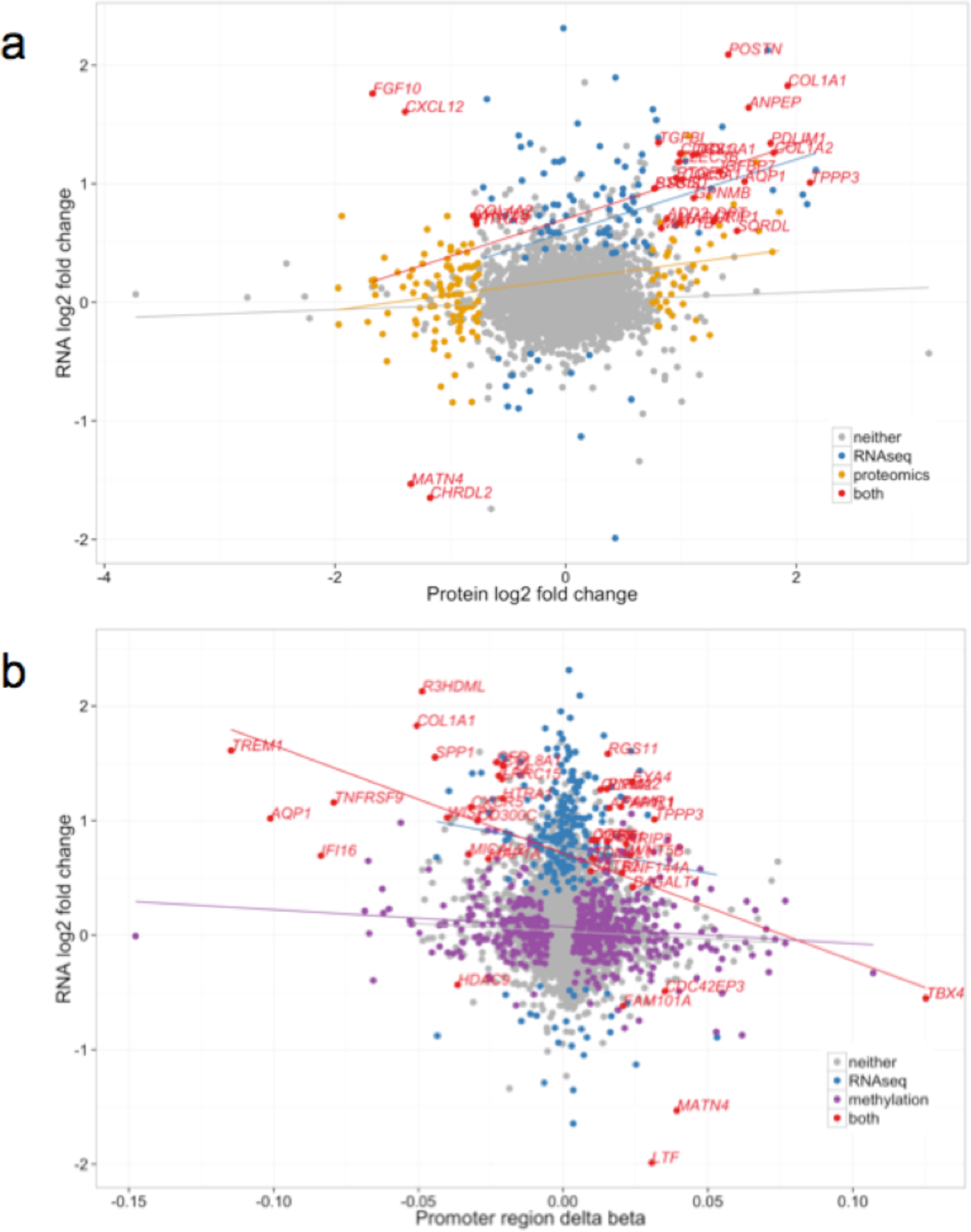
Comparison of changes identified in the ‐omics experiments. a) Comparison of the log-fold-changes between all genes identified in both the proteomics and RNA-seq experiments. Each gene is represented as a single point, and the colour corresponds to whether the gene is identified as differentially expressed using edgeR in the RNA-seq or proteomics experiments, or both. The trend lines are derived from a linear regression in each subset. Positive fold changes indicate increased expression in degraded samples. b) Comparison of RNA-seq log-fold-change, and mean promoter region methylation change. The trend lines are derived from a linear regression in each subset. Genes are coloured according to the results of the RNA-seq and the promoter-region analyses analogously to Figure 2a.

### RNA sequencing

We sequenced total RNA from all samples and identified 349 genes differentially expressed at a 5% false discovery rate (FDR). Of these, 296 and 53 genes demonstrated higher transcription levels in the degraded and intact samples, respectively (Supplementary Table 2). One of the most strongly down-regulated genes in chondrocytes isolated from damaged sites was *CHRDL2*, the presence of which was further confirmed by immunohistochemistry (IHC; Supplementary Figure 4). This gene was also found at lower abundance in the proteomics data.

### DNA methylation

We used the Illumina 450k methylation array to assay ~480,000 CpG sites across the genome. We first performed a probe level analysis and identified 9,896 differentially methylated probes (DMPs) at 5% FDR (Supplementary Table 3). We also identified 271 differentially methylated regions (DMRs), composed of multiple differentially methylated CpG sites (see **Methods**), associated with 296 unique overlapping genes (Supplementary Table 4).

### Integrative analyses: proteomics and RNA sequencing

Among the 209 proteins with evidence of differential abundance in the proteomics data, 31 were also differentially expressed at the RNA level (hypergeometric p=5.3x10^−7^, Figure 1b). Twenty-six of these 31 genes showed concordant directions of effect between degraded and intact samples (binomial p=0.0002), while the direction differed for five (*COL4A2*, *CXCL12*, *FGF10*, *HTRA3* and *WNT5B*). In all five cases the gene was found to be over-expressed at the RNA level and less abundant at the protein level in the degraded tissue. Based on the Human Protein Atlas [14], all five proteins encoded by these genes are annotated as predicted secreted proteins. In agreement with this, several collagens are more abundantly released into the culture media from diseased tissue than from healthy tissue [15].

We computed the global correlation in all samples irrespective of tissue status, comparing the RNA fragments per kilobase of transcript per million fragments mapped (FPKM) [16] to normalized peptide spectral counts (Supplementary Figure 5a) and found a significant positive correlation (Spearman’s rho=0.29, p<2.2x10^−16^) between RNA expression levels and protein abundance. To establish if there were also concordant differences in RNA and protein abundance between intact and degraded tissue, we computed the correlations between RNA and protein changes in degraded compared to intact samples (Figure 2a), and identified a significant positive correlation (Pearson’s r=0.17, p<2.2x10^−16^). The magnitude of correlation became substantially stronger when we only considered the 31 genes that were expressed differentially in both datasets (Pearson’s r=0.43, p=0.01).

### Integrative analyses: methylation and RNA sequencing

Sixteen of the genes with an associated DMR were also differentially transcribed (Figure 1b). For the direct comparison of methylation and gene expression in the following, we used the aggregate methylation status of promoter region CpG probes with transcription levels in all samples irrespective of intact/degraded status (see **Methods**, Supplementary Table 5). We found the expected negative correlation between promoter region methylation and gene expression (Spearman’s rho=-0.43, p<2.2x10^−16^, Supplementary Figure 5b). Based on the comparison of intact to degraded cartilage, the log-fold-changes in RNA expression and the differences in mean promoter region methylation values demonstrated a small but highly significant correlation (Pearson’s r=-0.08, p<2.2x10^−16^, Figure 2b). Again, the correlation became substantially higher when we considered the 39 genes with significant differences at both the promoter methylome and transciptome levels (Pearson’s r=-0.48, p=0.002).

### Integrative analyses: methylation, RNA sequencing, and proteomics

We identified 49 genes with evidence of differential regulation from at least two of the three–omics analyses (using the DMR methylation data; Supplementary Table 6), and three genes with consistent significant evidence for involvement in OA progression across all three levels: *AQP1*, *COL1A1* and *CLEC3B* (Figure 1b). All three genes were up-regulated in degraded tissue in both the RNA-seq and proteomics analyses (Figure 2a). *AQP1* and *COL1A1* showed a consistent decrease in methylation of all CpG probes in their associated DMRs, commensurate with an increase in transcription, while the DMR associated with *CLEC3B* showed evidence of increased methylation. Using IHC we independently confirmed the presence of *AQP1*, *COL1A1* and *CLEC3B* in articular cartilage chondrocytes (Supplementary Figure 4). We also replicated the direction of gene expression change for all three genes in independent data (see below and Supplementary Table 7).

Of the 49 genes with evidence of differential regulation on at least two molecular levels, 33 add substantive evidence to genes previously reported and 16 genes (33%) have not previously been implicated in OA (Supplementary Table 6).

### Replication of gene expression changes

We assayed gene expression in degraded and intact cartilage samples from two independent cohorts: a set of 17 patients with knee OA and a set of 9 patients with hip OA, using the same approach as for the discovery data (see **Methods**). After quality control, we retained 14,762 genes common to the discovery and both replication datasets, including 332 of 349 genes with FDR≤5% in the discovery data. We found excellent concordance in the direction of change for the genes with FDR≤5% in the discovery data: 93.4% of genes showed the same direction of effect in the knee replication data and 91.0% of genes showed the same direction of effect in the hip replication data (Figure 3a-b; both binomial p<10^−15^). Of the genes with concordant effect between the discovery and replication data, 65.5% reached nominal statistical significance in the knee replication data and 47% in the hip replication data (Supplementary Table 8; both binomial p<10^−15^).

**Figure 3:**
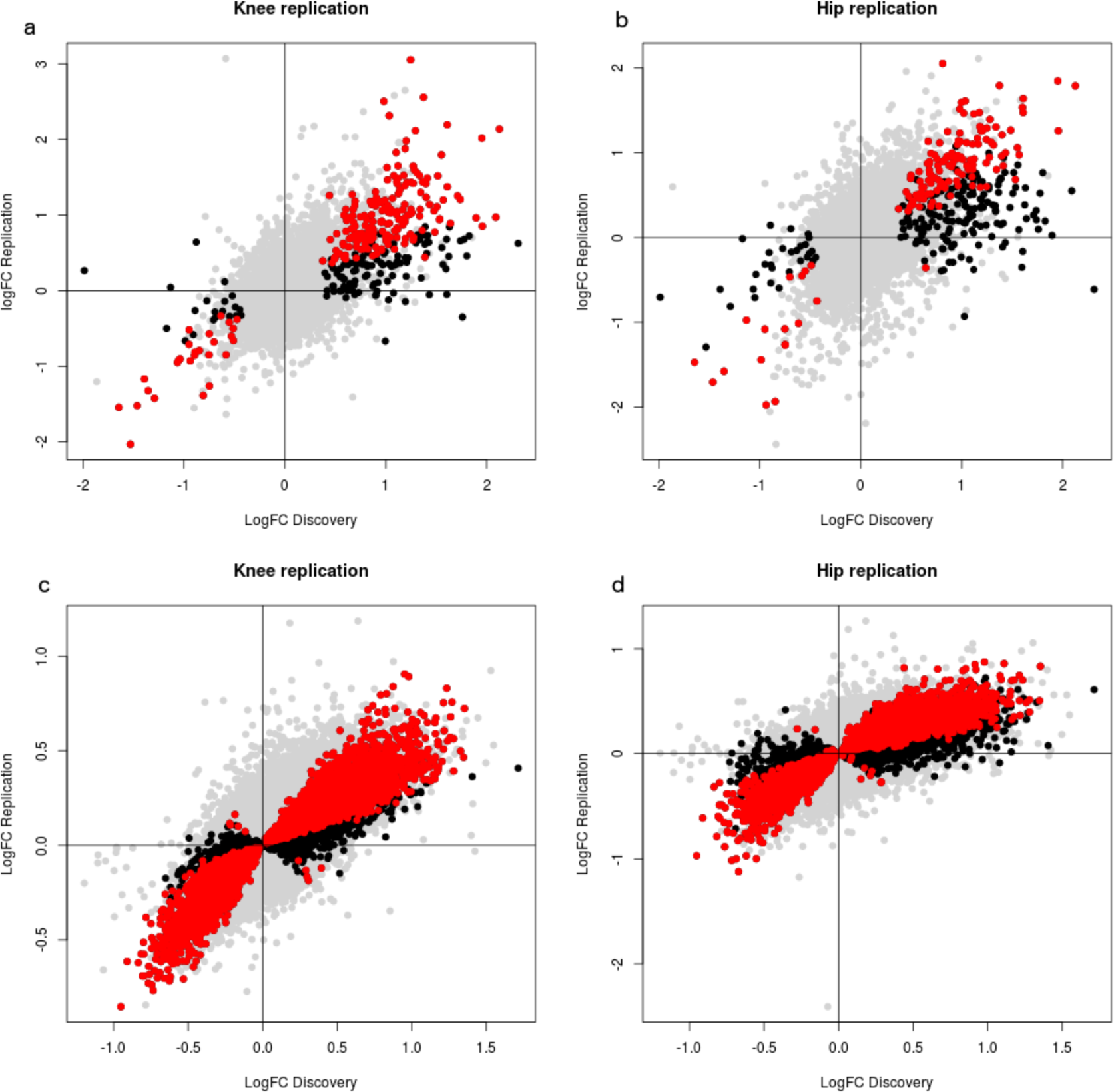
Replication of gene expression and methylation changes. a, b) Replication of gene expression changes in independent RNA-seq datasets of samples from patients with knee (a) and hip (b) OA. c, d) Replication of CpG methylation changes in independent datasets of samples from patients with knee (c) and hip (d) OA. Genes or probes with significant change at 5% in the discovery data are marked black; genes or probes that additionally show nominal significance in the replication data are marked in red.

Moreover, we found good correlation for the estimates of the log-fold-changes across all 14,762 genes between the knee discovery and the replication data: r=0.56 for knee replication data and r=0.51 for hip replication data (both p<10^−15^). These correlations are higher for the 332 genes with FDR≤5% in the discovery gene expression data (r=0.73 for knee, r=0.66 for hip replication data, both p<10^−15^). This shows that the gene expression changes identified in this study are robust and largely joint-independent.

We specifically considered the 49 genes with evidence from at least two ‐omics levels. Of these, 47 had gene expression data in the discovery and both replication datasets; 36 genes had nominally significant differential gene expression in the same direction in the knee replication data, and 26 genes in the hip replication data (Supplementary Table 7). This included *ANPEP*, for which we have used Western blotting to confirm protein changes (Supplementary Figure 3), and *CHRDL2*, for which we used immunohistochemistry to confirm presence of the protein (Supplementary Figure 4). Notably, the direction of change replicates in at least one of the knee and hip replication datasets at nominal significance for 13 of the 16 genes that have not previously been associated with OA (Supplementary Table 7).

We also pursued replication in an independent published microarray gene expression dataset of degraded and intact cartilage from the RAAK study, including 22 individuals with hip OA and 11 individuals with knee OA [17]. Of the 349 genes with FDR≤5% in the discovery data, 154 genes had expression measurements in the RAAK knee and hip replication datasets. We found highly significant agreement between the discovery and RAAK data: 83.8% of the genes showed the same direction of effect in the knee RAAK data (binomial one-sided p<10^−15^) and 69.5% of genes showed the same direction of effect in the hip RAAK data (binomial one-sided p<10^−6^). Furthermore, despite the difference in genomics technology (RNA-seq in discovery, microarray in RAAK), we found good concordance for the estimates of the log-fold-changes between the knee discovery and the RAAK replication data: r=0.43 for knee (p=3.6x10^−8^) and r=0.24 for hip replication data (p=0.003).

### Replication of methylation changes

To replicate the methylation results, we assayed DNA methylation in degraded and intact cartilage samples from two independent datasets: a set of 17 patients with knee OA and a set of 8 patients with hip OA, using the same approach as for the discovery data (see **Methods**). After quality control, we retained 416,437 probes common to the discovery and both replication data sets, including 9,723 of 9,867 differentially methylated probes (DMPs) with FDR≤5% in the knee discovery data. We found excellent concordance in the direction of change for the DMPs: 96.9% of probes showed the same direction of effect in the knee replication data and 95.2% probes showed the same direction of effect in the hip replication data (Figure 3c-d; both binomial one-sided p<10^−15^).

Furthermore, we found good correlation for the estimates of the fold-changes across all 416,437 probes between the knee discovery and the replication data: r=0.69 for knee replication data and r=0.56 for hip replication data (both p<10^−15^). The correlation is even higher for the DMPs: r=0.91 for knee, r=0.85 for hip replication data (both p<10^−15^). Similarly to the gene expression data, this shows that the methylation changes estimated in this study are robust and largely joint-independent.

In summary, our combined epigenetic, transcriptomic and proteomic analysis has uncovered a substantial number of genes associated with OA progression, some of which have known connections to cartilage or bone-related processes, and others with no links reported to date, bringing new potential insights into the molecular mechanisms of OA pathogenesis. The replication data confirm the high quality of our discovery experiment and further support a large number of the novel genes with a suggested role in OA progression.

### Gene set analyses

We performed a gene set enrichment analysis on the genes with significant evidence for differential expression, methylation and/or protein abundance from each separate–omics analysis and found that several common biological processes are highlighted at multiple levels (Supplementary Tables 9-10). We additionally identified pathways jointly affected by genes identified at multiple molecular levels (Figure 4, Supplementary Figure 6, Supplementary Tables 9-10).

**Figure 4:**
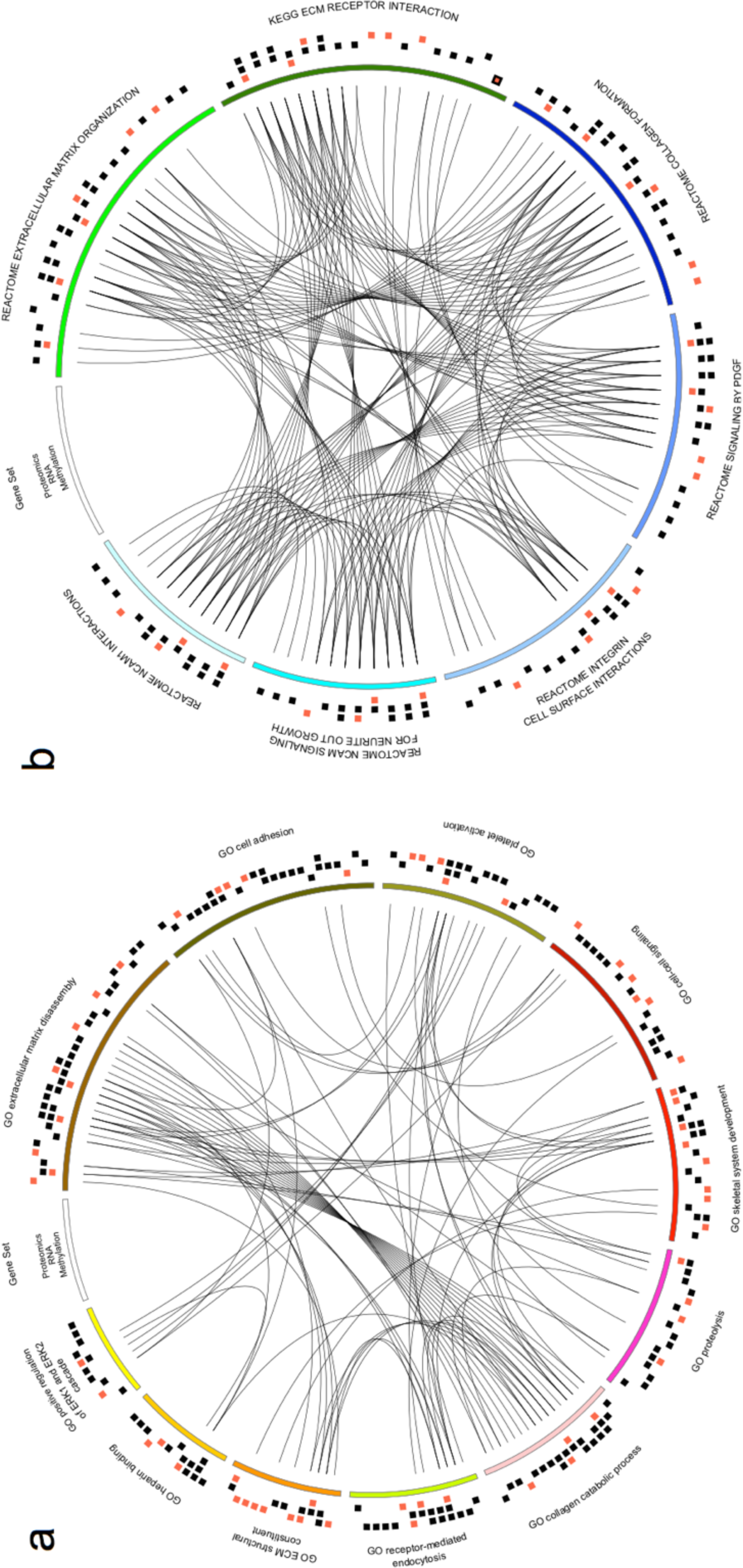
Significant gene set enrichments in the integrative ‐omics analysis. a, b) Enrichments from KEGG/Reactome (a) and Gene Ontology (b). The circos plots show enriched gene sets, with genes differentially regulated in at least one of the methylation, RNA-seq, or proteomics experiments. Lines connect genes that occur in several gene sets. The three outside circles show boxes for genes with significantly higher (black) or lower (red) methylation, gene, or protein expression data. A red box with black border indicated a gene that overlaps hyper‐ as well as hypo-methylated DMRs.

A strong theme in the highlighted pathways is cartilage matrix regulation and degeneration, confirming the fact that increased ECM turnover is a crucial component in OA pathogenesis. Pathways including extracellular matrix organization and collagen formation were also implicated by genes identified by all three ‐omics analyses (Figure 4). Results from the three analyses converge on shared mechanisms, supporting the importance of taking an integrated perspective. The GO term analysis also uncovered consistent evidence from all three ‐omics assays for genes annotated for extracellular matrix disassembly and collagen catabolic process. In these pathways we also find suggestive evidence of a link to genetic OA risk loci (see Supplementary Methods and Results, Supplementary Table 11). These signals would not necessarily have been identified directly from GWAS data, highlighting the importance of synthesizing information from multiple molecular levels to obtain a more powerful integrated view.

Positive regulation of ERK1/2 cascade, heparin-binding and platelet activation were also enriched at multiple molecular levels and are interconnected through common genes. Several studies have linked the extracellular signal-regulated kinase (ERK) cascade to OA [5, 18-21]. Heparin-binding growth factors have also been shown to be involved in OA [22-25], some in particular through activation of the ERK signaling pathway. Injection of platelet-rich plasma in OA knees leads to significant clinical improvement [26, 27] and there is evidence to suggest that this effect is mediated via the ERK cascade [28]. Our findings provide strong evidence supporting a role for this pathway in OA pathogenesis.

We also found enrichment of genes involved in the regulation of angiogenesis at multiple levels. The growth of blood vessels and nerves are closely linked processes that share regulatory mechanisms, including the ERK cascade and heparin-binding proteins mentioned above [29]. Accordingly, NCAM signaling for neurite outgrowth and PDGF signaling that play a significant role in blood vessel and nervous system formation were highlighted by the pathway analysis. We found significant enrichment in plasma proteins for both the RNA-seq (hypergeometric p=6.9x10^−11^) and the proteomics experiments (p=1.8x10^−5^). This supports a role for angiogenesis and nerve growth in OA progression [29, 30]. Indeed, histological examination of the samples we investigated showed greater blood vessel ingrowth in tissues with more advanced OA (Supplementary Figure 1). Results from the three molecular analyses converge on shared biological mechanisms that are relevant to the pathogenesis of OA. These data should be useful in pinpointing candidate targets to help improve therapeutic intervention.

### *In silico* screen for drug targets

To identify existing drugs that could be applied to OA, we searched Drugbank [31] using the 49 differentially regulated genes identified by at least two of the functional genomics approaches. We uncovered 29 compounds with investigational or established actions on the corresponding proteins. After filtering to include only agents with current Food and Drug Administration Marketing Authorization for use in humans, we identified ten agents with actions on nine of the dysregulated proteins (see Supplementary Results, Supplementary Table 12). For the corresponding nine genes, eight have RNA-seq data, and the direction of change replicates for seven genes in the knee and six genes in the hip RNA-seq replication data at nominal significance (Supplementary Tables 8,12). Consequently, this work could help prioritize the repurposing of existing drugs for the treatment of OA.

## DISCUSSION

Previous studies of osteoarthritis have investigated methylation [32, 33], transcription [17, 34, 35], or protein expression [36, 37] in OA tissue separately, and others have combined up to two of these ‐omics assays [38, 39], some with the addition of genetic data [40]. By contrast, this study provides the first integrated, systematic and hypothesis-free analysis of the biological changes involved in human cartilage OA progression using genome-wide data at three molecular levels. Using this multi-level functional genomics approach, we have provided a first integrated view of the molecular alterations within chondrocytes that accompany cartilage degeneration leading to debilitating end-stage joint disease. All data arising from the experiments described here are freely available to the wider scientific community, from the raw data to the analysis results for each ‐omics level.

We show that the direction of effect for over 90% of gene expression and CpG methylation changes replicate in independent knee or hip OA data. We also replicate the majority of gene expression changes using independent knee and hip OA data from a microarray experiment. This supports the quality of our data and suggests that there are largely shared molecular changes in knee and hip OA progression, within the power constraints of our study.

We have focused on improving our understanding of the molecular processes that underpin disease progression and have used matched intact and degraded cartilage samples from knee OA joints. Our data highlight 49 genes with evidence of differential regulation at multiple molecular levels, identifying several novel genes, including *MAP1A* and *PXDN*, and providing robust cross-cutting evidence for genes previously implicated in OA, including *AQP1*, *COL1A1,* and *CLEC3B*.

*AQP1* encodes aquaporin-1, a member of a family of proteins that facilitate water transport across biological membranes. Chondrocyte swelling and increased cartilage hydration has been suggested as an important mechanism in OA [41]. Accordingly, *AQP1* has been observed to be over-expressed in a rat model of knee OA [42], and in knee OA in humans [43].

*CLEC3B* (also known as *TNA*) encodes the protein tetranectin, which binds human tissue plasminogen activator (tPA) [44]. Previous studies have identified *CLEC3B* as up-regulated in human OA [34, 45], mediating extracellular matrix destruction in cartilage and bone [46], and a candidate gene association study found evidence of association of a coding variant (rs13963, Gly106Ser) in *CLEC3B* with OA [47] (although this association has not been replicated in subsequent studies [48]).

*COL1A1* is one of several collagen proteins, which were differentially abundant at both the RNA-seq and proteomics levels (including *COL1A2*, *COL3A1*, *COL4A2*, *COL5A1*, Supplementary Table 6). Collagens are the main structural components of cartilage and several studies have highlighted the importance of collagen dysregulation in OA [6, 49]. A recent study also identified up-regulation of *COL1A1* and *COL5A1* in synovium from humans with end-stage OA, in the synovium of mice with induced OA and in human fibroblasts stimulated with TGF-β [50].

*MAP1A* and *MAP1B,* two novel genes with convergent evidence confirmed by replication, encode microtubule-associated proteins, both of which were up-regulated significantly at both the RNA and protein levels (Figure 2a). These proteins are expressed mostly in the brain and are involved in regulation of the neural cytoskeleton [51]. Cytoskeletal regulation is thought to be an important process in OA [52] and accordingly, recent studies have implicated these proteins in bone formation [53].

The *PXDN* gene was up-regulated at the RNA level, confirmed by replication, and associated with 2 hypo-methylated DMRs. *PXDN* encodes peroxidasin, which is secreted into the extracellular matrix and catalyses collagen IV cross-linking [54]. The other replicated genes not previously implicated in OA (Supplementary Table 6) have relatively little characterization.

Among the genes we identify that have been previously implicated in OA, *ANPEP* (aminopeptidase E) is a broad specificity aminopeptidase that has previously been detected in the synovial fluid of OA patients [55] and therefore has potential as a novel OA biomarker. *CHRDL2* is a bone morphogenetic protein (*BMP*) inhibitor that has previously been reported to be lost from chondrocytes of the superficial zone and shifted to the middle zone in OA cartilage in a targeted study [56]. *WNT5B*, a ligand for frizzled receptors in the WNT signaling pathway, has previously been reported to be differentially transcribed in osteoarthritic bone, consistent with current understanding that OA is a disease involving both cartilage and bone [57].

On the level of gene sets, we identify extracellular matrix organization, collagen catabolism and angiogenesis as biological pathways that are clearly implicated in disease progression.

Intervention to prevent disease progression in OA as well as targeted prevention in high-risk disease-free individuals will be important in reducing the future societal burden of OA [58]. A clearer understanding of the factors that modulate OA disease progression is thus critical to the development of novel prognostic markers and biological disease-modifying agents, analogous to those emerging or successfully applied in inflammatory arthritis [59-61]. Follow-up mechanistic studies will be required before causal relationships between the identified pathways and OA progression can be established.

In the chondrocyte, we found little evidence of differential inflammatory pathway activity between the intact and degraded samples [62, 63]. This is not surprising, as all of the patients whose tissues were studied here had a diagnosis of OA and clinically advanced disease in at least 1 location within the joint. Inflammatory mediators are soluble factors present throughout the joint, to which both the healthy and the diseased chondrocyte populations are potentially exposed, and for which the regulatory molecules may differ to those prioritized by the tissues studied here.

Based on our discovery data from 12 individuals, we estimate that ~10% of the “true” differentially expressed genes are statistically significant in this study and ~95% of significantly different genes are true positives (Supplementary Figure 7). The high true positive rate is also confirmed by the replication data for gene expression and methylation.

Larger sample sizes will be required for a more powerful characterization of the molecular changes occurring with disease progression. The integrative functional genomics approach illustrated here offers an opportunity to identify molecular signatures in disease-relevant tissues, thereby gaining insights into disease mechanism, identifying potential biomarkers, and discovering druggable targets for intervention. A key future challenge will be the development of powerful statistical approaches for the integration of high-dimensional molecular traits in the context of complex diseases.

## METHODS

### Patient consent & study approval

All subjects provided written, informed consent prior to participation in the study. Tissue samples were collected under Human Tissue Authority license 12182, Sheffield Musculoskeletal Biobank, University of Sheffield, UK. All samples were collected from patients undergoing total knee replacement for primary osteoarthritis. The patients comprised 2 women and 10 men, mean age 66 years (range 50-88). Patients with diagnosis other than osteoarthritis were excluded from the study. The study was approved by Oxford NHS REC C (10/H0606/20).

### Sample processing

*Extraction of chondrocytes from osteochondral tissue taken at knee replacement* Osteochondral samples were transported in Dulbecco’s modified Eagle’s medium (details see Supplementary Methods). Half of each sample was taken for chondrocyte extraction and the remaining tissue was fixed in 10% neutral buffered formalin, decalcified in surgipath decalcifier (Leica) and embedded to paraffin wax for histological and immunohistochemical analysis. Chondrocytes were directly extracted from each paired macroscopic intact and degraded OA cartilage sample in order to remove the extracellular matrix allowing a higher yield of cells to be loaded onto the Qiagen column.

#### Histological examination

Four micron sections of paraffin-embedded cartilage tissue were mounted onto positively charged slides and histologically stained using Haematoxylin and Eosin, Alcian blue, Masson trichrome (details see Supplementary Methods). Cartilage tissue was graded using the Mankin Score (0-14) with additional scores for abnormal features (0-4) and cartilage thickness (0-4) based on the OARSI scoring system [12, 13]. The total scores were used to determine the overall grade of the cartilage as healthy/low-grade degenerate, referred to as “intact” (median: 4.5; IOR: 3-5.5; n=12), or high-grade degenerate, referred to as “degraded” (median: 14; IOR: 11.75-18; n=12).

#### Extraction of DNA, RNA, and protein

Detailed protocols are included in the Supplementary Methods. In brief, cartilage was removed from the bone, dissected and washed twice in 1xPBS. Tissue was digested overnight, passed through a cell strainer, centrifuged, washed twice, and re-suspended. Cells were counted using a haemocytometer and the viability checked using trypan blue exclusion (Invitrogen). The optimal cell number for spin column extraction from cells was between 4x106 and 1x107. Cells were then pelleted and homogenized. DNA, RNA and protein extractions were performed using the Qiagen AllPrep DNA/RNA/Protein Mini Kit, as per manufacturer’s instructions. RNA, DNA and protein were quantitated by picogreen and gel electrophoresis. Samples were frozen at -80 degrees C prior to assays.

### Proteomics

Detailed protocols for protein digestion, TMT labelling, peptide fractionation and all individual steps below are included in the Supplementary Methods.

#### LC-MS Analysis

LC-MS analysis was performed on the Dionex Ultimate 3000 UHPLC system coupled with the high-resolution LTQ Orbitrap Velos mass spectrometer (Thermo Scientific).

The ten most abundant multiply charged precursors within 380 -1500 *m/z* were selected with FT mass resolution of 30,000 and isolated for HCD fragmentation with isolation width 1.2 Th. Tandem mass spectra were acquired with FT resolution of 7,500 and targeted precursors were dynamically excluded for further isolation and activation for 40 seconds with 10 ppm mass tolerance. Samples from one individual were excluded during quality control.

#### Database Search and Protein Quantification

The acquired mass spectra were submitted to SequestHT search engine implemented on the Proteome Discoverer 1.4 software for protein identification and quantification, with settings as described in the Supplementary Methods. Peptide confidence was estimated with the Percolator node. Peptide FDR was set at 0.01 and validation was based on q-value and decoy database search. All spectra were searched against a UniProt fasta file containing 20,190 Human reviewed entries. The Reporter Ion Quantifier node included a custom TMT 6plex Quantification Method with integration window tolerance 20 ppm and integration method the Most Confident Centroid. For each identified protein a normalized spectral count value was calculated for each one of the 6-plex experiments by dividing the number of peptide spectrum matches (PSMs) of each protein with the total number of PSMs. Median normalized spectral counts per protein were computed across the different multiplex experiments.

#### Differential abundance

To identify those proteins with evidence of differential expression, we shortlisted proteins with absolute median abundance ratios between degraded and intact samples ≥0.75, where the median abundance ratio was greater than the standard deviation in all samples with data, and with evidence from at least 5 patients. This analysis identified 209 proteins (Supplementary Table 1).

#### Western blotting

Sample pairs were adjusted to the same protein concentration. Primary antibodies used were as follows: ANPEP, ab108382; AQP1, ab168387; COL1A, ab14918; TGFB1, ab89062; WNT5B, ab124818 (Abcam); GAPDH, sc-25778 (Santa Cruz Biotechnologies). Intensity values were normalized to GAPDH loading control before ratio calculation. Further details are in the Supplementary Methods.

#### Label free quantification of representative samples

For a selection of four representative control and disease samples, peptide aliquots of 500ng without TMT labelling were analyzed on the Dionex Ultimate 3000 UHPLC system coupled with the Orbitrap Fusion (Thermo Scientific) mass spectrometer for label free quantification and validation, as described in the Supplementary Methods. With a minimum requirement of at least total 14 spectra per protein we found excellent agreement in the direction of change between isobaric labelling and label free quantification for at least 32 proteins which is approximately 90% of the common proteins between the TMT changing list and the label free identified list (Supplementary Figure 2).

### RNA-seq

Detailed protocols for all individual steps are included in the Supplementary Methods.

#### RNA sequencing

The steps for mRNA purification, cDNA library creation, and litigation to Illumina Paired-end Sequencing adaptors are described in the Supplementary Methods. The libraries then went through 10 cycles of PCR amplification using KAPA Hifi Polymerase rather than the kit-supplied Illumina PCR Polymerase due to better performance.

Samples were quantified and pooled based on a post-PCR Agilent Bioanalyzer, then the pool was size-selected using the LabChip XT Caliper. The multiplexed library was then sequenced on the Illumina HiSeq 2000, 75bp paired-end read length. Sequenced data was then analyzed and quality controlled (QC) and individual indexed library BAM files were produced.

#### Read alignment

The resulting reads that passed QC were realigned to the GRCh37 assembly of the human genome using a splice-aware aligner, bowtie version 2.2.3 [64], and using a reference transcriptome from Ensembl release 75 [65], using the ‐library-type option fr-firststrand to bowtie. We limited the alignments to uniquely mapping reads. We then counted the number of reads aligning to each gene in the reference transcriptome using htseq-count from the HTSeq package [66] separately for each sample to produce a read count matrix counting the number of reads mapping to each gene in the transcriptome for each sample. To quantify absolute transcript abundance we computed the fragments per kilobase of transcript per million fragments mapped (FPKM) [16] for each gene using the total read counts from this matrix, and the exonic length of each gene calculated from gene models from Ensembl release 75. We obtained a mean of 49.3 million uniquely mapping reads from each sample (range: 39.2-71.4 million) with a mean of 84% of reads mapping to genes (range: 67.9- 90.6%) which were used for the differential expression analysis.

#### Differential expression analysis

We used edgeR version 3.0 [67] to identify differentially expressed genes from the read count matrix. We restricted the analysis to 15,418 genes with >1 counts per million in at least 3 samples (similar to the protocol described by [68]). We followed the processing steps listed in the manual, using a generalized linear model with tissue status (degraded or intact) and individual ID as covariates. 349 genes were differentially expressed between the degraded and intact samples at 5% FDR (296 up-, 54 down-regulated in degraded tissue). The genes differentially expressed at 5% FDR had somewhat higher exonic length than the remaining genes (Wilcox-test *p*=0.00013; 4804 vs 4153 bases), hence we adjusted for gene length in the randomizations for gene set analyses (see Supplementary Methods).

### Methylation

We used the Illumina 450k BeadChip to assay methylation, with sample preparation (including Bisulfite Conversion, pre‐ and post-amplification) described in detail in the Supplementary Methods.

#### Illumina 450k BeadChip assay

BeadChips were scanned on five Illumina iScans, four of which are paired with two Illumina Autloader 2.Xs. The iScan software produced intensity files for each channel (.idat files).

#### Probe-level analysis

The intensity files for each sample were processed using the ChAMP package [69]. Probes mapping to chromosomes X & Y, and those with a detection p-value >0.01 (n=3,064) were excluded. The beta values for each probe were quantile-normalized, accounting for the design of the array, using the ‘dasen’ method from the wateRmelon package [70]. We also excluded any probes with a common SNP (minor allele frequency >5%) within 2 base pairs of the CpG site, and those predicted to map to multiple locations in the genome [71] (n=45,218), leaving a total of 425,694 probes for the probe-level differential methylation analysis. We annotated all probes with genomic position, gene and genic location information from the ChAMP package.

To identify probes with evidence of differential methylation we used the CpGassoc package [72] to fit a linear model at each probe, with tissue status and individual ID as covariates. This analysis yielded 9,867 differentially methylated probes (DMP) between degraded and intact samples at 5% FDR.

#### Region-level analysis

To identify differentially methylated regions, we used custom software (available upon request) to identify regions containing at least 3 DMPs and no more than 3 non-significant probes with no more than 1kb between each constituent probe, following previous analyses [39]. We used bedtools [73] to identify genes overlapping each DMR, using gene annotations from Ensembl release 75, and extending each gene’s bound to include 1500 basepairs upstream of the transcription start site to include likely promoter regions. This analysis yielded 271 DMRs with a mean of 4.04 DMPs per region, and a mean length of 673 basepairs.

#### Promoter-level analysis

We assigned probes in the promoter region of each gene using the probe annotations from the ChAMP package, and assigned to each gene any probe with the annotation “TSS1500”, “TSS200”, “5’UTR” and “1stExon” in order to capture probes in likely promoter regions. We then computed the mean normalized beta value of assigned probes for all genes with at least 5 associated probes for each sample separately, to produce a single methylation value for each gene in each sample. We used a paired t-test to identify genes with differential promoter-region methylation between degraded and intact samples, and a 5% FDR cutoff to call a gene’s promoter region as differentially methylated. Note that the paired t-test assumes an equivalent model to the linear model used for the probe-level analysis.

### Immunohistochemistry

To identify whether native chondrocytes demonstrated expression of the key factors immunohistochemistry was deployed. The detailed protocol is included in the Supplementary Methods.

### Protein atlas annotation

We downloaded annotations from Human Protein Atlas version 13, and annotated each protein-coding gene from the 3 experiments with the following terms taken from the annotation file: “Predicted secreted protein”, “Predicted membrane protein”, “Plasma protein”. The secreted and membrane protein predictions are based on a consensus call from multiple computational prediction algorithms, and the plasma protein annotations are taken from the Plasma Protein Database, as detailed in [14].

### Identification of previously reported OA genes

In order to identify whether some of the genes we highlight have previously been reported as associated with OA, we searched PubMed on 2 September 2016. We used an “advanced” search of the form “(osteoarthritis) AND (<gene_name>)” where <gene_name> was set to each HGNC gene symbol and we report the number of citations returned for each search.

### Replication of gene expression changes using RNA-seq data and replication of methylation changes

### Knee samples

Tissue samples were collected under National Research Ethics approval reference 15/SC/0132, South Yorkshire and North Derbyshire Musculoskeletal Biobank, University of Sheffield. All samples were collected from patients undergoing total knee replacement for primary osteoarthritis, and all patients provided written informed consent before participation. The patients comprised 12 women and 5 men, mean age 71 years (range 54- 82). Patients with diagnosis other than osteoarthritis were excluded from the study. All sample processing steps (extraction of chondrocytes, extraction of DNA) were carried out as for the knee OA discovery samples.

### Hip samples

Tissue samples were collected under national Research Ethics approval reference 11/EE/0011, Cambridge Biomedical Research Centre Human Research Tissue Bank, Cambridge University Hospitals, UK. All samples were collected from patients undergoing total hip replacement for primary osteoarthritis. The patients, who provided written consent before participation, comprised 6 women and 3 men, mean age 61 years (range 44-84). Patients with diagnosis other than osteoarthritis were excluded from the study. All sample processing steps (extraction of chondrocytes, extraction of DNA) were carried out as for the knee OA discovery samples, except that the cartilage was digested overnight in 6mg/ml collagenase A (Roche) in medium containing 10% serum to release the cells.

### Replication analysis for gene expression changes: RNA-seq data

We applied the same procedure to the knee and hip replication data as to the discovery data. We considered 14762 genes that passed QC in the knee discovery, knee replication, and hip replication data; this included 332 of 349 genes with FDR<=5% in the knee discovery data.

### Replication analysis for methylation changes

We used the Illumina 450k BeadChip to assay methylation for all knee and hip replication samples, using the same procedure as for the knee discovery data. The knee and hip methylation data were processed using the same QC procedure as for the knee discovery samples, including the same QC thresholds. We excluded one degraded hip cartilage sample as an outlier in probe failure rate (over six times the proportion of the next highest sample), and also excluded the paired intact cartilage sample. After QC, 417077 probes remained. We considered 416437 probes with methylation values in the knee discovery, knee replication, and hip replication data; this included 9723 of 9867 probes with FDR<=5% in the knee discovery data (“DMPs”).

### Replication of gene expression changes: microarray data

OA-dependent changes in expression of genes with differential expression in the discovery data were assessed with the help of an available dataset from the ongoing Research Arthritis and Articular Cartilage (RAAK) study, consisting in gene expression profiles of OA affected cartilage and macroscopically preserved cartilage from 33 patients undergoing total joint replacement surgery (22 with hip OA, 11 with knee OA). Sample collection and determination of gene expression levels have been described in detail previously [17]. In short, cartilage was collected separately for the OA affected and the unaffected regions of the weight baring part of the joint, snap frozen in liquid nitrogen and stored at -80°C prior to RNA extraction. Gene expression was determined with the Illumina HumanHT-12 v3 microarrays. After removal of probes that were not optimally measured (detection p-value>0.05 in more than 50% of the samples) a paired t-test was performed on all sample pairs while adjusting for chip (to adjust for possible batch effects).

### Gene set analyses

#### Individual datasets

We aimed to test whether particular biological gene sets were enriched among the significant genes from each of the RNA-seq, methylation, and proteomics datasets. To this end, we downloaded KEGG [74] and Reactome [75] gene annotations from MSigDB (version 4) [76]. We also downloaded Gene Ontology (GO) biological process and molecular function gene annotations from QuickGO [77] on 4 February 2015. For GO, we only considered annotations with evidence codes IMP, IPI, IDA, IEP, and TAS. Genes annotated to the same term were treated as a “pathway”. KEGG/Reactome and GO annotations were analysed separately and only pathways with 20 to 200 genes were considered (555 for KEGG/Reactome, 677 for GO). Enrichment was assessed using a 1-sided hypergeometric test and only considering genes with annotations from a particular resource, as described in the Supplementary Methods. Multiple-testing was accounted for by using a 5% FDR (separately for KEGG/Reactome and GO, and for RNA-seq, methylation, and protein expression data).

Empirical p-values for the enrichments were obtained from randomizations accounting for overlap of significant genes among the RNA-seq, methylation, and protein expression datasets (see Supplementary Methods).

#### Integrative gene set analyses

We aimed to integrate the gene sets analyses for the RNA-seq, methylation, and protein expression datasets. For each gene set, we asked whether the association across the three datasets (calculated as geometric mean of the *p*-values) was higher than expected by chance. To this end, we obtained 1-sided empirical *p*-values from 100,000 sets of “random RNA-seq genes, random methylation genes, and random protein expression genes”. The “random” sets were chosen to conservatively match the overlap observed among the significant genes, and to account for gene length as described in the Supplementary Methods. We confirmed that the number of randomizations was sufficient (see Supplementary Methods). We performed the randomization separately for KEGG/Reactome and for GO, as we only considered genes with at least one annotation in the resource.

Analyses using the arcOGEN genetic data are described in the Supplementary Methods and in the Supplementary Results.

## DECLARATIONS

### Ethics approval and consent to participate

See Methods.

### Consent for publication

Not applicable.

### Availability of data and material

All genome-wide summary statistics generated in the discovery component of this study are available as Supplementary Information. All discovery RNA sequencing data are available in EGA repository, https://www.ebi.ac.uk/ega/studies/EGAS00001001203. All discovery methylation data are available in EGA repository, https://www.ebi.ac.uk/ega/studies/EGAS00001001213, respectively. The discovery proteomics data has been deposited in the PRIDE archive under study ID PXD002014 [publication in process].

## Acknowledgements

The authors wish to thank Sara Dunn and Clive Buckle for contribution to extraction of RNA/DNA/protein from chondrocyte samples, Danielle Walker for research administration, and Pei-Chien Tsai and Jordana Bell for advice on the methylation analyses. The authors would like to acknowledge the contribution of the Wellcome Trust Sanger Institute Sample Management, Illumina Bespoke, and Genotyping teams to this work. This study utilized genotype data from arcOGEN (http://www.arcogen.org.uk/) funded by a special purpose grant from Arthritis Research UK (grant 18030). This study makes use of data generated by the Wellcome Trust Case-Control Consortium (the 1958 British Birth Cohort collection and the UK Blood Services Collection). A full list of the investigators who contributed to the generation of the data is available from www.wtccc.org.uk. We thank all participants of the RAAK study. We would like to acknowledge the Cambridge Biomedical Research Center Human Research Tissue Bank for storing the hip gene expression and methylation replication samples.

## Authors’ contributions

Study design: EZ, JMW. Omics data analysis: JS, GRSR, TIR. Discovery data sample collection: JMW. Knee replication data (non-RAAK) sample collection: RLJ, KMS, JMW. Hip replication data (non-RAAK) sample collection (including protocol development): RAB, AWM. Proteomics: JSC, TIR. Histology and immunohistochemistry: CLLM, ALAB. Western blotting: MP, RC. Gene expression replication data from RAAK study: YFMR, RGHHN, IM. Manuscript writing and editing: JS, GRSR, TIR, RAB, MP, CLLM, YFMR, IM, AWM, JSC, JMW, EZ.

## REFERENCES

1. Vos T, Flaxman AD, Naghavi M, Lozano R, Michaud C, Ezzati M, Shibuya K, Salomon JA, Abdalla S, Aboyans V, et al: Years lived with disability (YLDs) for 1160 sequelae of 289 diseases and injuries 1990–2010: a systematic analysis for the Global Burden of Disease Study 2010. The Lancet 2012, 380:2163-2196.

2. Dieppe PA, Lohmander LS: Pathogenesis and management of pain in osteoarthritis. The Lancet 2005, 365:965-973.

3. Valdes AM, Spector TD: Genetic epidemiology of hip and knee osteoarthritis. Nat Rev Rheumatol 2011, 7:23-32.

4. Lawrence RC, Helmick CG, Arnett FC, Deyo RA, Felson DT, Giannini EH, Heyse SP, Hirsch R, Hochberg MC, Hunder GG, et al: Estimates of the prevalence of arthritis and selected musculoskeletal disorders in the United States. Arthritis & Rheumatism 1998, 41:778-799.

5. Lee AS, Ellman MB, Yan D, Kroin JS, Cole BJ, van Wijnen AJ, Im H-J: A current review of molecular mechanisms regarding osteoarthritis and pain. Gene 2013, 527:440-447.

6. Xia B, Di C, Zhang J, Hu S, Jin H, Tong P: Osteoarthritis Pathogenesis: A Review of Molecular Mechanisms. Calcified Tissue International 2014, 95:495-505.

7. Yuan XL, Meng HY, Wang YC, Peng J, Guo QY, Wang AY, Lu SB: Bone– cartilage interface crosstalk in osteoarthritis: potential pathways and future therapeutic strategies. Osteoarthritis and Cartilage 2014, 22:1077-1089.

8. Reynard LN, Loughlin J: The genetics and functional analysis of primary osteoarthritis susceptibility. Expert Rev Mol Med 2013, 15.

9. Ruiz-Romero C, Fernández-Puente P, Calamia V, Blanco FJ: Lessons from the proteomic study of osteoarthritis. Expert Review of Proteomics 2015, 12:433-443.

10. Ramos YF, Meulenbelt I: The role of epigenetics in osteoarthritis: current perspective. Curr Opin Rheumatol 2016.

11. Steinberg J, Zeggini E: Functional genomics in osteoarthritis: Past, present, and future. J Orthop Res 2016, 34:1105-1110.

12. Mankin HJ, Dorfman H, Lippiello L, Zarins A: Biochemical and metabolic abnormalities in articular cartilage from osteo-arthritic human hips. II. Correlation of morphology with biochemical and metabolic data. J Bone Joint Surg Am 1971, 53:523-537.

13. Pearson RG, Kurien T, Shu KSS, Scammell BE: Histopathology grading systems for characterisation of human knee osteoarthritis – reproducibility, variability, reliability, correlation, and validity. Osteoarthritis and Cartilage 2011, 19:324-331.

14. Uhlen M, Fagerberg L, Hallstrom BM, Lindskog C, Oksvold P, Mardinoglu A, Sivertsson A, Kampf C, Sjostedt E, Asplund A, et al: Tissue-based map of the human proteome. Science 2015, 347:1260419-1260419.

15. Lourido L, Calamia V, Mateos J, Fernández-Puente P, Fernández-Tajes J, Blanco FJ, Ruiz-Romero C: Quantitative Proteomic Profiling of Human Articular Cartilage Degradation in Osteoarthritis. J Proteome Res 2014, 13:6096-6106.

16. Trapnell C, Williams BA, Pertea G, Mortazavi A, Kwan G, van Baren MJ, Salzberg SL, Wold BJ, Pachter L: Transcript assembly and quantification by RNA-Seq reveals unannotated transcripts and isoform switching during cell differentiation. Nat Biotechnol 2010, 28:511-515.

17. Ramos YFM, den Hollander W, Bovée JVMG, Bomer N, van der Breggen R, Lakenberg N, Keurentjes JC, Goeman JJ, Slagboom PE, Nelissen RGHH, et al: Genes Involved in the Osteoarthritis Process Identified through Genome Wide Expression Analysis in Articular Cartilage; the RAAK Study. PLoS ONE 2014, 9:e103056.

18. Liu Z, Cai H, Zheng X, Zhang B, Xia C: The Involvement of Mutual Inhibition of ERK and mTOR in PLC?1-Mediated MMP-13 Expression in Human Osteoarthritis Chondrocytes. IJMS 2015, 16:17857-17869.

19. Otero M, Plumb DA, Tsuchimochi K, Dragomir CL, Hashimoto K, Peng H, Olivotto E, Bevilacqua M, Tan L, Yang Z, et al: E74-like Factor 3 (ELF3) Impacts on Matrix Metalloproteinase 13 (MMP13) Transcriptional Control in Articular Chondrocytes under Proinflammatory Stress. Journal of Biological Chemistry 2012, 287:3559-3572.

20. Prasadam I, Zhou Y, Shi W, Crawford R, Xiao Y: Role of dentin matrix protein 1 in cartilage redifferentiation and osteoarthritis. Rheumatology 2014, 53:2280-2287.

21. Xu J, Yi Y, Li L, Zhang W, Wang J: Osteopontin induces vascular endothelial growth factor expression in articular cartilage through PI3K/AKT and ERK1/2 signaling. Mol Med Rep 2015, 12:4708-4712.

22. Chia S-L, Sawaji Y, Burleigh A, McLean C, Inglis J, Saklatvala J, Vincent T: Fibroblast growth factor 2 is an intrinsic chondroprotective agent that suppresses ADAMTS-5 and delays cartilage degradation in murine osteoarthritis. Arthritis Rheum 2009, 60:2019-2027.

23. Long DL, Ulici V, Chubinskaya S, Loeser RF: Heparin-binding epidermal growth factor-like growth factor (HB-EGF) is increased in osteoarthritis and regulates chondrocyte catabolic and anabolic activities. Osteoarthritis and Cartilage 2015, 23:1523-1531.

24. Patil AS, Sable RB, Kothari RM: Occurrence, biochemical profile of vascular endothelial growth factor (VEGF) isoforms and their functions in endochondral ossification. J Cell Physiol 2012, 227:1298-1308.

25. Pufe T, Groth G, Goldring MB, Tillmann B, Mentlein R: Effects of pleiotrophin, a heparin-binding growth factor, on human primary and immortalized chondrocytes. Osteoarthritis and Cartilage 2007, 15:155-162.

26. Marmotti A, Rossi R, Castoldi F, Roveda E, Michielon G, Peretti GM: PRP and Articular Cartilage: A Clinical Update. BioMed Research International 2015, 2015:1-19.

27. Meheux CJ, McCulloch PC, Lintner DM, Varner KE, Harris JD: Efficacy of Intra-articular Platelet-Rich Plasma Injections in Knee Osteoarthritis: A Systematic Review. Arthroscopy: The Journal of Arthroscopic & Related Surgery 2016, 32:495-505.

28. Zhou Q, Xu C, Cheng X, Liu Y, Yue M, Hu M, Luo D, Niu Y, Ouyang H, Ji J, Hu H: Platelets promote cartilage repair and chondrocyte proliferation via ADP in a rodent model of osteoarthritis. Platelets 2015, 27:212-222.

29. Mapp PI, Walsh DA: Mechanisms and targets of angiogenesis and nerve growth in osteoarthritis. Nat Rev Rheumatol 2012, 8:390-398.

30. Ashraf S, Walsh DA: Angiogenesis in osteoarthritis. Current Opinion in Rheumatology 2008, 20:573-580.

31. Law V, Knox C, Djoumbou Y, Jewison T, Guo AC, Liu Y, Maciejewski A, Arndt D, Wilson M, Neveu V, et al: DrugBank 4.0: shedding new light on drug metabolism. Nucleic Acids Research 2013, 42:D1091-D1097.

32. Jeffries MA, Donica M, Baker LW, Stevenson ME, Annan AC, Humphrey MB, James JA, Sawalha AH: Genome-Wide DNA Methylation Study Identifies Significant Epigenomic Changes in Osteoarthritic Cartilage. Arthritis & Rheumatology 2014, 66:2804-2815.

33. Moazedi-Fuerst FC, Hofner M, Gruber G, Weinhaeusel A, Stradner MH, Angerer H, Peischler D, Lohberger B, Glehr M, Leithner A, et al: Epigenetic differences in human cartilage between mild and severe OA. Journal of Orthopaedic Research 2014, 32:1636-1645.

34. Karlsson C, Dehne T, Lindahl A, Brittberg M, Pruss A, Sittinger M, Ringe J: Genome-wide expression profiling reveals new candidate genes associated with osteoarthritis. Osteoarthritis and Cartilage 2010, 18:581-592.

35. Tew SR, McDermott BT, Fentem RB, Peffers MJ, Clegg PD: Transcriptome-Wide Analysis of Messenger RNA Decay in Normal and Osteoarthritic Human Articular Chondrocytes. Arthritis & Rheumatology 2014, 66:3052-3061.

36. Stenberg J, Rüetschi U, Skiöldebrand E, Kärrholm J, Lindahl A: Quantitative proteomics reveals regulatory differences in the chondrocyte secretome from human medial and lateral femoral condyles in osteoarthritic patients. Proteome Sci 2013, 11:43.

37. Ruiz-Romero C, Carreira V, Rego I, Remeseiro S, López-Armada MJ, Blanco FJ: Proteomic analysis of human osteoarthritic chondrocytes reveals protein changes in stress and glycolysis. Proteomics 2008, 8:495-507.

38. den Hollander W, Ramos YFM, Bomer N, Elzinga S, van der Breggen R, Lakenberg N, de Dijcker WJ, Suchiman HED, Duijnisveld BJ, Houwing‐Duistermaat JJ, et al: Transcriptional Associations of Osteoarthritis‐ Mediated Loss of Epigenetic Control in Articular Cartilage. Arthritis & Rheumatology 2015, 67:2108-2116.

39. den Hollander W, Ramos YFM, Bos SD, Bomer N, van der Breggen R, Lakenberg N, de Dijcker WJ, Duijnisveld BJ, Slagboom PE, Nelissen RGHH, Meulenbelt I: Knee and hip articular cartilage have distinct epigenomic landscapes: implications for future cartilage regeneration approaches. Annals of the Rheumatic Diseases 2014, 73:2208-2212.

40. Rushton MD, Reynard LN, Young DA, Shepherd C, Aubourg G, Gee F, Darlay R, Deehan D, Cordell HJ, Loughlin J: Methylation quantitative trait locus analysis of osteoarthritis links epigenetics with genetic risk. Human Molecular Genetics 2015, 24:7432-7444.

41. Bush PG, Hall AC: The volume and morphology of chondrocytes within non-degenerate and degenerate human articular cartilage. Osteoarthritis and Cartilage 2003, 11:242-251.

42. Musumeci G, Leonardi R, Carnazza ML, Cardile V, Pichler K, Weinberg AM, Loreto C: Aquaporin 1 (AQP1) expression in experimentally induced osteoarthritic knee menisci: An in vivo and in vitro study. Tissue and Cell 2013, 45:145-152.

43. Geyer M, Grässel S, Straub RH, Schett G, Dinser R, Grifka J, Gay S, Neumann E, Müller-Ladner U: Differential transcriptome analysis of intraarticular lesional vs intact cartilage reveals new candidate genes in osteoarthritis pathophysiology. Osteoarthritis and Cartilage 2009, 17:328-335.

44. Westergaard UB, Andersen MH, Heegaard CW, Fedosov SN, Petersen TE: Tetranectin binds hepatocyte growth factor and tissue-type plasminogen activator. Eur J Biochem 2003, 270:1850-1854.

45. Valdes AM, Hart DJ, Jones KA, Surdulescu G, Swarbrick P, Doyle DV, Schafer AJ, Spector TD: Association study of candidate genes for the prevalence and progression of knee osteoarthritis. Arthritis & Rheumatism 2004, 50:2497-2507.

46. Herren T: Regulation of plasminogen receptors. Frontiers in Bioscience 2003, 8:d1-8.

47. Valdes AM, Van Oene M, Hart DJ, Surdulescu GL, Loughlin J, Doherty M, Spector TD: Reproducible genetic associations between candidate genes and clinical knee osteoarthritis in men and women. Arthritis Rheum 2006, 54:533-539.

48. Panoutsopoulou K, Zeggini E: Advances in osteoarthritis genetics. Journal of Medical Genetics 2013, 50:715-724.

49. Tchetina EV: Developmental Mechanisms in Articular Cartilage Degradation in Osteoarthritis. Arthritis 2011, 2011:1-16.

50. Remst DFG, Blom AB, Vitters EL, Bank RA, van den Berg WB, Blaney Davidson EN, van der Kraan PM: Gene Expression Analysis of Murine and Human Osteoarthritis Synovium Reveals Elevation of Transforming Growth Factor β-Responsive Genes in Osteoarthritis‐Related Fibrosis. Arthritis & Rheumatology 2014, 66:647-656.

51. Halpain S, Dehmelt L: The MAP1 family of microtubule-associated proteins. Genome Biology 2006, 7:224.

52. Blain EJ: Involvement of the cytoskeletal elements in articular cartilage homeostasis and pathology. International Journal of Experimental Pathology 2009, 90:1-15.

53. Kanenari M, Zhao J, Abiko Y: Enhancement of microtubule-associated protein-1 Alpha gene expression in osteoblasts by low level laser irradiation. Laser Ther 2011, 20:47-51.

54. Péterfi Z, Geiszt M: Peroxidasins: novel players in tissue genesis. Trends in Biochemical Sciences 2014, 39:305-307.

55. Balakrishnan L, Nirujogi R, Ahmad S, Bhattacharjee M, Manda SS, Renuse S, Kelkar DS, Subbannayya Y, Raju R, Goel R, et al: Proteomic analysis of human osteoarthritis synovial fluid. Clin Proteomics 2014, 11:6.

56. Nakayama N: A novel chordin-like BMP inhibitor, CHL2, expressed preferentially in chondrocytes of developing cartilage and osteoarthritic joint cartilage. Development 2004, 131:229-240.

57. Hopwood B, Tsykin A, Findlay DM, Fazzalari NL: Microarray gene expression profiling of osteoarthritic bone suggests altered bone remodelling, WNT and transforming growth factor-β/bone morphogenic protein signalling. Arthritis Res Ther 2007, 9:R100.

58. Hiligsmann M, Cooper C, Guillemin F, Hochberg MC, Tugwell P, Arden N, Berenbaum F, Boers M, Boonen A, Branco JC, et al: A reference case for economic evaluations in osteoarthritis: An expert consensus article from the European Society for Clinical and Economic Aspects of Osteoporosis and Osteoarthritis (ESCEO). Seminars in Arthritis and Rheumatism 2014, 44:271-282.

59. Goulielmos GN, Zervou MI, Myrthianou E, Burska A, Niewold TB, Ponchel F: Genetic data: The new challenge of personalized medicine, insights for rheumatoid arthritis patients. Gene 2016, 583:90-101.

60. Mease PJ: Biologic Therapy for Psoriatic Arthritis. Rheumatic Disease Clinics of North America 2015, 41:723-738.

61. Wood AJJ, Olsen NJ, Stein CM: New Drugs for Rheumatoid Arthritis. New England Journal of Medicine 2004, 350:2167-2179.

62. Goldring MB, Otero M: Inflammation in osteoarthritis. Current Opinion in Rheumatology 2011, 23:471-478.

63. Wojdasiewicz P, Poniatowski ŁA, Szukiewicz D: The Role of Inflammatory and Anti-Inflammatory Cytokines in the Pathogenesis of Osteoarthritis. Mediators of Inflammation 2014, 2014:1-19.

64. Langmead B, Salzberg SL: Fast gapped-read alignment with Bowtie 2. Nature Methods 2012, 9:357-359.

65. Flicek P, Amode MR, Barrell D, Beal K, Billis K, Brent S, Carvalho-Silva D, Clapham P, Coates G, Fitzgerald S, et al: Ensembl 2014. Nucleic Acids Research 2013, 42:D749-D755.

66. Anders S, Pyl PT, Huber W: HTSeq—a Python framework to work with high-throughput sequencing data. Bioinformatics 2014, 31:166-169.

67. Robinson MD, McCarthy DJ, Smyth GK: edgeR: a Bioconductor package for differential expression analysis of digital gene expression data. Bioinformatics 2009, 26:139-140.

68. Anders S, McCarthy DJ, Chen Y, Okoniewski M, Smyth GK, Huber W, Robinson MD: Count-based differential expression analysis of RNA sequencing data using R and Bioconductor. Nat Protoc 2013, 8:1765-1786.

69. Morris TJ, Butcher LM, Feber A, Teschendorff AE, Chakravarthy AR, Wojdacz TK, Beck S: ChAMP: 450k Chip Analysis Methylation Pipeline. Bioinformatics 2014, 30:428-430.

70. Pidsley R, Y Wong CC, Volta M, Lunnon K, Mill J, Schalkwyk LC: A datadriven approach to preprocessing Illumina 450K methylation array data. BMC Genomics 2013, 14:293.

71. Chen Y-a, Lemire M, Choufani S, Butcher DT, Grafodatskaya D, Zanke BW, Gallinger S, Hudson TJ, Weksberg R: Discovery of cross-reactive probes and polymorphic CpGs in the Illumina Infinium HumanMethylation450 microarray. Epigenetics 2013, 8:203-209.

72. Barfield RT, Kilaru V, Smith AK, Conneely KN: CpGassoc: an R function for analysis of DNA methylation microarray data. Bioinformatics 2012, 28:1280-1281.

73. Quinlan AR, Hall IM: BEDTools: a flexible suite of utilities for comparing genomic features. Bioinformatics 2010, 26:841-842.

74. Kanehisa M: KEGG: Kyoto Encyclopedia of Genes and Genomes. Nucleic Acids Research 2000, 28:27-30.

75. Matthews L, Gopinath G, Gillespie M, Caudy M, Croft D, de Bono B, Garapati P, Hemish J, Hermjakob H, Jassal B, et al: Reactome knowledgebase of human biological pathways and processes. Nucleic Acids Research 2009, 37:D619-D622.

76. Subramanian A, Tamayo P, Mootha VK, Mukherjee S, Ebert BL, Gillette MA, Paulovich A, Pomeroy SL, Golub TR, Lander ES, Mesirov JP: Gene set enrichment analysis: A knowledge-based approach for interpreting genome-wide expression profiles. Proceedings of the National Academy of Sciences 2005, 102:15545-15550.

77. Binns D, Dimmer E, Huntley R, Barrell D, O’Donovan C, Apweiler R: QuickGO: a web-based tool for Gene Ontology searching. Bioinformatics 2009, 25:3045-3046.

